# Potential genetic robustness of *Prnp* and *Sprn* double knockout mouse embryos towards ShRNA-lentiviral inoculation

**DOI:** 10.1101/2021.10.22.465458

**Authors:** Andrea Rau, Bruno Passet, Johan Castille, Alexandre Asset, Jérome Lecardonnel, Marco Moroldo, Florence Jaffrézic, Denis Laloë, Katayoun Moazami-Goudarzi, Jean-Luc Vilotte

## Abstract

Shadoo, encoded by *Sprn*, and PrP, encoded by *Prnp*, are related proteins whose biological functions are still incompletely understood. Although previous knockdown experiments have suggested the necessity of Shadoo in the absence of PrP during early mouse embryogenesis, little impact of the double-knockout of these two loci was reported. To further investigate this apparent discrepancy, we compared the transcriptome of WT, *Prnp*^*0/0*^ and *Prnp*^*0/0*^, *Sprn*^*0/0*^ E6.5 mouse embryos following inoculation by *Sprn*-ShRNA or *Prnp*-ShRNA lentiviral vectors at the one-cell stage. Our results highlighted a significant induction of an apoptotic pathway in *Prnp*^*0/0*^ E6.5 mouse embryos inoculated with *Sprn*-ShRNA vectors alongside interferon and to a lesser extent inflammatory responses, confirming previous reported experiments. On the contrary, ShRNA vector inoculation in *Prnp*^*0/0*^, *Sprn*^*0/0*^ embryos did not induce apoptosis and resulted in lower interferon responses. Finally, comparisons of the transcriptome of WT and *Prnp*^*0/0*^, *Sprn*^*0/0*^ embryos revealed only slight differences, which may in part explain the genetic robustness observed in the latter genotype.

**Highlights:**

- *Sprn-ShRNA* lentivirus vector inoculation in *Prnp* knockout one-cell mouse embryos results in the induction of an apoptosis pathway at E6.5, alongside interferon and to a lesser extent inflammatory responses.
- *Sprn- or Prnp-ShRNA* lentivirus vector inoculations in *Prnp*/*Sprn* knockout one-cell mouse embryos induce lower interferon responses and no apoptotic pathway at E6.5.
- Although wild type and *Prnp*/*Sprn* knockout E6.5 mouse embryos are transcriptomically similar, some differences might explain this apparent resilience of the double knockout genotype.

## 1. Introduction

PrP, the prion protein encoded by *Prnp* whose abnormally folded isoform is the key component of infectious prion particles responsible for Transmissible Spongiform Encephalopathies, and Shadoo, encoded by *Sprn*, are evolutionarily related [1]. Their biological functions remain incompletely understood, and their individual genetic invalidations result in no overt phenotypes beyond resistance to prion infection for *Prnp*-knockout mice. The early embryonic lethality, already noticeable at E7.5 with a developmental failure of the trophectoderm-derived compartment, of *Prnp*-knockout, *Sprn*-knockdown embryos initially suggested a biological redundancy of these two proteins [2]. However, the development of *Prnp* and *Sprn* double knockout mice in various genetic backgrounds [3,4] did not confirm this hypothesis. These apparently contradictory observations could result from a genetic compensation in invalidated animals [5] or from an increased robustness [6].

In the present report, we comparatively assessed, at the transcriptomic level, the impact of *Prnp* and *Sprn* knockout in E6.5 mouse embryos and its consequences following inoculation with *ShRNA*-lentiviral vectors at the one cell stage.

## 2. Materials and Methods

### 2.1 Ethics statements

Animal experiments were carried out in accordance with the EU Directive 2010/63/EU. Production, breeding and inoculations at the one cell stage with *ShRNA*-lentiviral vectors of the various transgenic lines were approved by the Ethics Committee of Jouy-en-Josas (Comethea, Permit number 02532.01), and followed the safety recommendations of the French “Haut Conseil des Biotechnologies” (HCB, Permit numbers 6460 and 5468).

### 2.2 Transgenic lines and lentiviral inoculations

Transgenic *Prnp* and *Sprn* FVB/N knockout mouse lines were already described [2,4,7]. Wild type (WT) FVB/N mice were purchased from Janvier (https://www.janvier-labs.com/fiche_produit/souris_fvb). ShhRNA lentiviral vector solutions were purchased from Sigma with infectious titers over 10^9^ infectious units/ml (LS1: TRCN0000179960 and LS2: TRCN0000184740 against *Sprn* transcripts, LP1: TRCN0000319687 and LP2: TRCN 0000273801 against *Prnp* transcripts). Intra-perivitellin space injections and transplantation into pseudo-pregnant recipient mice were performed as previously described [2]. Around 50 one-cell stage embryos were injected for each genotype and lentiviral solution combination (Figure 1A).

**Figure 1:**
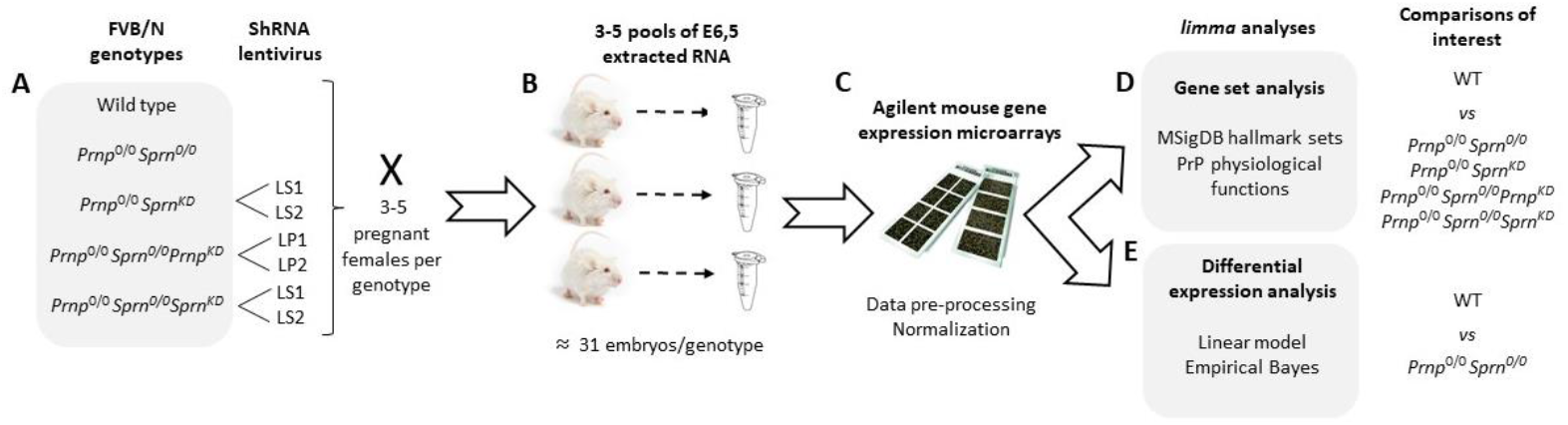
Schematic representation of study design and analysis. (A) Transgenic lines and lentiviral inoculations; (B) embryo collection; (C) transcriptomic analyses; (D) hallmark gene set analysis; and (E) differential expression analysis.

### 2.3 Embryo collection and transcriptomic analyses

Embryos were collected at E6.5 (Figure 1B). Total RNA was isolated from pools of 6-14 E6.5 embryos, deriving from 3 to 5 females. RNA extractions and integrity analysis were performed as previously described [2]. Three to four independent pools were produced for each experimental group (i.e., genotype and lentiviral solution combination) and analysed using Agilent SurePrint G3 gene expression V2 8×60K mouse microarrays (AMADID: 074809, Figure 1C). All steps were performed by the @BRIDGe facility (INRAE Jouy-en-Josas, France, http://abridge.inra.fr/), as described previously [8].

All analyses were performed with R version 4.0.0. Median pixel intensity and local background intensity were read and pre-processed from the raw Agilent files using the R/Bioconductor package *limma* (version 3.44.1, [9,10]). Probe intensities were quantile-normalized and log_2_-transformed [11]. Using the “gIsWellAboveBackground” flag, non-control probes were called as present if they were above background in at least 3 samples. After averaging intensities for remaining probes with identical target sequences, a single representative probe was chosen for each gene according to the maximum observed variance across samples (Figure 1C). Raw microarray data files and all analysis scripts needed to pre-process the data and reproduce the analyses described in this work are openly available on the Data INRAE portal at https://data.inrae.fr/privateurl.xhtml?token=e7b8885d-9cef-43c9-ae98-06259ee84d44.

### 2.4 Hallmark gene set analyses

To evaluate the potential role played by specific ensembles of gene sets, hallmark gene sets from the Molecular Signatures Database (MSigDB, [13,14]) were obtained for *Mus musculus* using the *msigdbr* package (version 7.2.1). Among the 50 available gene sets, we focused our attention on a subset of 14 hallmark gene sets related to PrP recognized physiological functions (see below). Comparisons of interest for the hallmark gene set analysis were defined for four different experimental groups as compared to WT mice: (1) *Prnp*^*0/0*^, *Sprn*^*0/0*^; (2) *Prnp*^0/0^, *Sprn*^*0/0*^, *Sprn*^*KD*^; (3) *Prnp*^0/0^, *Sprn*^*0/0*^, *Prnp*^*KD*^; and (4) *Prnp*^0/0^, *Sprn*^*KD*^. To minimize possible off-target effects, contrasts for comparisons with groups (2)-(4) were constructed by averaging over the two lentiviruses for each gene knockdown. Using the *fry* self-contained rotation gene set test from *limma* [15], we sought to identify whether genes in each selected hallmark gene set were globally differentially expressed for a given comparison (Figure 1D). *P*-values were calculated corresponding to tests for gene sets exhibiting significant over-expression (“Up”) and under-expression (“Down”), as well as differential expression regardless of direction (“Mixed”). Raw *P*-values were corrected for multiple testing using the Benjamini-Hochberg approach to control the false discovery rate (FDR, [12]), and gene sets were identified as significantly globally differentially expressed if their adjusted *P-*value < 0.05.

### 2.5 Differential expression analysis

For the differential analysis (Figure 1E), a linear model with group-means parameterization (i.e., no intercept and a separate coefficient for each group) was fit for each gene. Using *limma*, an empirical Bayes approach was used to moderate the standard errors of the estimated log-fold changes. Contrasts were defined to identify differentially expressed genes for each comparison of interest; we focused in particular on the comparison of *Prnp*^*0/0*^, *Sprn*^*0/0*^ and WT E6.5 embryos. As before, *P-*values were corrected for multiple testing using the Benjamini-Hochberg approach to control the false discovery rate [13], and genes were identified as significantly differentially expressed if their adjusted *P*-value < 0.05 and absolute log fold change > 1.

## 3. Results and discussion

### 3.1 Knockdown of Sprn in mouse Prnp^0/0^ embryos induces apoptosis alongside interferon responses at E6.5

Although *Prnp*^*0/0*^,*Sprn*^*0/0*^ mice are viable [3,4], the knockdown of *Sprn* in *Prnp*^*0/0*^ mouse embryos was reported to induce embryonic lethality highlighted by a developmental failure of the trophectoderm-derived compartment noticeable at E7.5 [2]. We reinvestigated this latter observation by transcriptomic analysis of such embryos at E6.5, focusing on a subset of MSigDB including 14 hallmark gene sets related to PrP recognized physiological functions ([1,16-18], Table 1). Three of those hallmark gene sets were significantly altered in *Sprn*-knockdown, *Prnp*^*0/0*^ E6.5 embryos compared to their WT counterparts (adjusted P-value < 0.05): interferon-α and –γ responses and apoptosis, while inflammatory response was significant with an adjusted P-value < 0.10 (Table 1). These data confirm the above-mentioned previous report, emphasizing that embryonic lethality could be diagnosed at earlier developmental stages. Because two different ShRNA were used, targeting different regions of the *Sprn* transcript, it is unlikely that apoptosis results from an off-target effect. We next wondered if this apoptotic induction, alongside inflammatory and interferon responses, could result from the association of a lentiviral ShRNA-expressing vector inoculation [19] with a *Prnp*-knockout induced interferon-primed state [20] in the absence/reduction of *Sprn* expression, as it was previously shown that this latter is required to induce apoptosis [2].

**Table 1:**
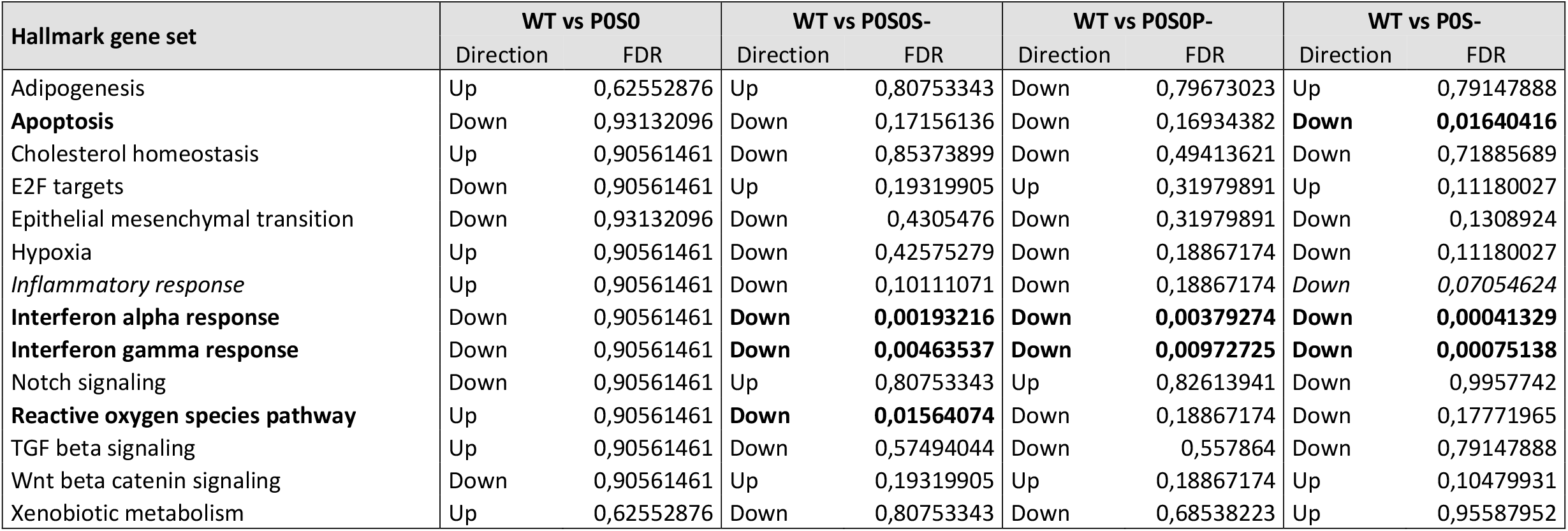
Hallmark gene set analyses at E6.5. Top margin: Compared genotypes. P0: *Prnp*^*0/0*^. SO: *Sprn*^*0/0*^. S-: knockdown of *Sprn*. P-: knockdown of *Prnp*. For each knockdown, two independent lentiviral ShRNA vectors were used (see Material and Methods). Left margin: hallmark gene sets [13,14]. Significantly altered hallmark gene sets are highlighted in boldface (FDR < 0.05) and italicized (FDR < 0.10).

### 3.2 Prnp or Sprn knockdown in mouse Prnp^0/0^, Sprn^0/0^ embryos induces reduced interferon responses and no apoptosis at E6.5

We similarly investigated the transcriptomic outcomes at E6.5 of *Sprn*- or *Prnp*-knockdown in *Prnp*^*0/0*^, *Sprn*^*0/0*^ mouse embryos. The knockdown of *Sprn* or *Prnp* were performed on a knockout genotype for both these genes to highlight only those pathways associated with lentiviral ShRNA vector infections on this specific genetic background. Two different ShRNA were again used for each targeted gene to reduce the likelihood of observing an off-target-induced biological disturbance. Compared to WT E6.5 embryos, only two hallmark gene sets were altered consistently and significantly: interferon-α and –γ responses (Table 1). However, compared to the previous analysis, the statistical significance of these gene sets was unexpectedly reduced by 10-fold. Furthermore, no apoptosis induction was detected (Table 1).

These results could suggest that the expression or the knockout of *Sprn* is required to avoid lentiviral ShRNA vector induction of a strong interferon response associated with apoptosis in *Prnp*^*0/0*^ mouse embryos, while its knockdown exacerbates these pathways. A potential explanation for these apparent contradictory observations is that the knockout of the two genes induces a genetic adaptation that in turn helps control the lentiviral-induced responses. Such an adaptation might not take place with the *Sprn*-knockdown or to an insufficient level.

### 3.3 Transcriptomes of WT and Prnp^0/0^, Sprn^0/0^ E6.5 embryos are highly similar with only few differentially expressed genes

To assess this hypothesis, we next compared the transcriptome of E6.5 WT and *Prnp*^*0/0*^, *Sprn*^*0/0*^ mouse embryos. Only 11 genes were found to be differentially expressed between these two genotypes with an adjusted P-value < 0.05 and absolute log fold change > 1 (Table 2). All 11 of these genes were similarly found to be significantly differentially expressed in the same direction in *Sprn*-knockdown, *Prnp*^*0/0*^ compared to WT E6.5 mouse embryos, albeit with weaker log fold changes for the majority. As already discussed, the *Prnp* and *Sprn* gene invalidations did not induce alteration of their transcript expression levels and their absence in this list was thus expected [4,7]. Most of the differentially expressed genes were reported to be transcribed in the embryo ectoderm and mesenchyme, and only few in the endoderm or in the extraembryonic component (http://www.informatics.jax.org/expression.shtml, Table 2). Since adult expression of both *Prnp* and *Sprn* genes is more abundant in the nervous system, and since PrP involvement in muscle and bone development/regeneration has been previously reported, deregulation of these genes in the ectoderm and in the mesenchyme might be relevant observations. However, in *Sprn*-knockdown, *Prnp*^*0/0*^ embryos, a developmental failure of the trophectoderm-derived compartment was reported [2], rather suggesting a major role of the extraembryonic component in the appearance of this lethality.

**Table 2:** Differentially expressed genes between *Prnp*^*0/0*^, *Sprn*^*0/0*^ and WT E6.5 mouse embryos. Results are shown for significantly differentially expressed genes (FDR < 0.05, absolute log fold change ≥ 1). Checkmarks for each gene represent reported expression in embryo ectoderm, embryo endoderm, embryo mesenchyme, and extraexmbryonic component (http://www.informatics.jax.org/expression.shtml). Blank spaces and dashes represent unreported expression and no available data, respectively.

| Gene Name | Description | NCBI Gene | Log Fold Change | Adjusted P-Value | Embryo Ectoderm | Embryo Endoderm | Embryo Mesenchyme | Extraembryonic Component |
| --- | --- | --- | --- | --- | --- | --- | --- | --- |
| Spint1 | Serine protease inhibitor, Kunitz type 1 | 20732 | 1,599051 | 1,96414E-08 | ✓ | ✓ | ✓ | ✓ |
| Gm30906 | Long non-coding RNA | 102632964 | -1,477668 | 1,96414E-08 | ✓ |  |  |  |
| Scg5 | Secretogranin V, secreted chaperone protein | 20394 | -1,334427 | 4,2595E-08 | ✓ |  | ✓ |  |
| Spg11 | Spatacsin vesicle trafficking associated | 214585 | -1,081024 | 2,05026E-06 | ✓ |  | ✓ |  |
| Cds2 | CDP-diacylglycerol synthase 2 | 110911 | 1,297156 | 2,42507E-06 | ✓ |  | ✓ | ✓ |
| Jmjd7 | Jumonji domain containing 7 | 433466 | -1,479236 | 1,31005E-05 | ✓ |  | ✓ |  |
| Ada | Adenosine deaminase | 11486 | -2,117557 | 4,32324E-05 | ✓ |  | ✓ | ✓ |
| AK148702 |  | RIKEN clone 7120437D13 | 2,819679 | 0,004937838 | --- | --- | --- | --- |
| Cplx2 | Complexin 2 | 12890 | 2,616736 | 0,010457547 | ✓ |  | ✓ |  |
| Gm10734 | - | RIKEN clone I530011G18 | -1,104109 | 0,010457547 | ✓ |  | ✓ |  |
| Sdc4 | Syndecan 4 | 20971 | -1,157117 | 0,021156447 | ✓ |  | ✓ |  |

Only 3 out of the 11 genes are expressed in the extraembryonic component: *Spint1, Cds2* and *Ada* (Table 2). *Spint1* was recently reported to be a biomarker of placental insufficiency [21]. Low circulating levels of *Spint1* are associated with placental failure whereas here, at E6.5, this expression is higher in *Prnp*^*0/0*^, *Sprn*^*0/0*^ embryos compared to their WT counterparts. Whether *Spint1* overexpression can favor placental development remains to be demonstrated. *Cds2* is a widely expressed gene indirectly involved in the positive control of angiogenesis [22]. Its overexpression in *Prnp*^*0/0*^, *Sprn*^*0/0*^ embryos could suggest a sustained angiogenesis of the placenta, but in the absence of associated deregulation of co-factors, such as vascular endothelial growth factors, the interpretation of this observation remains fragile. Nevertheless, the differential expression of the two above-mentioned genes appears to favor placental development and to contribute to the survival of the *Prnp*^*0/0*^, *Sprn*^*0/0*^ mouse embryos. However, their potential implication in the control of the interferon response remains elusive.

### 3.4 Ada downregulation in Prnp^0/0^, Sprn^0/0^ E6.5 embryos could explain the apparent genotype robustness

The third gene, transcribed in the extraembryonic component and strongly differentially expressed (Log fold change −2.1, Table 2, Figure 2) is *Ada*. Disruption of the *Ada* gene in mice induces perinatal lethality [23], a phenotype rescued by tissue-specific placental expression of this gene [24]. Its crucial role in the trophectoderm-derived compartment was also indirectly emphasized through the knockout of the AP-2γ transcription factor-encoding gene that resulted in an early embryonic lethal phenotype, similar to that observed for *Sprn*-knockdown, in *Prnp*^*0/0*^ embryos [2,25], associated with a lack of *Ada* gene expression in the extraembryonic cells [26]. However, in *Prnp*^*0/0*^, *Sprn*^*0/0*^ mouse embryos, only a downregulation of the *Ada* gene expression is observed, thus likely avoiding the occurrence of these drastic phenotypes. It should be mentioned that *Ada*^*0/+*^ mouse embryos were similarly not reported to be affected [23].

**Figure 2:**
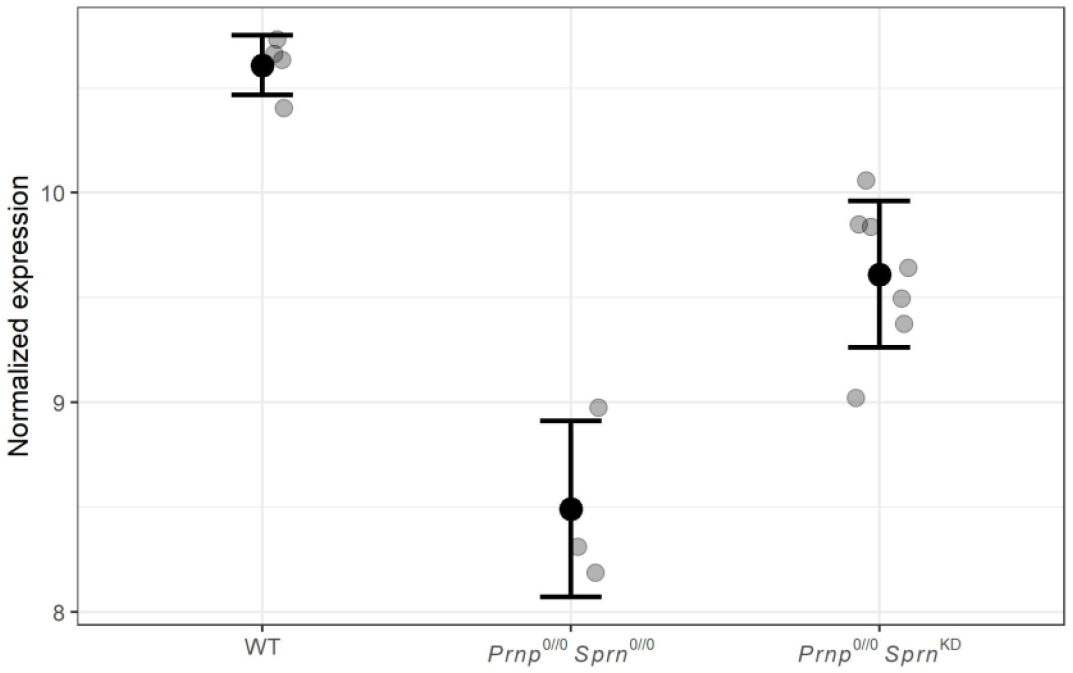
Normalized expression of *Ada* in WT; *Prnp*^*0/0*^, *Sprn*^*0/0*^; and *Prnp*^*0/0*^, *Sprn*^*KD*^ mice represented as dot plots for individual samples (grey points) with means (black points) and standard deviations (bars) for each experimental group.

Interestingly, Ada congenital defect induces a severe combined immunodeficiency syndrome [25]. Expression levels of this enzyme correlate with the production levels of interferons and proinflammatory factors, and modulation of Ada activity was even proposed as a potential therapeutic target [26-29]. High interferon responses can induce side effects among which some, such as autoimmune reactions, can be detrimental. The control of the interferon response is thus crucial, and as already mentioned, altered in the absence of members of the prion protein family [16]. The downregulation of the *Ada* gene expression observed in *Prnp*^*0/0*^, *Sprn*^*0/0*^ mouse embryos might help to control the interferon and inflammatory responses induced by lentiviral ShRNA-encoding vector infections to a level compatible with their survival. This genetic adaptation is only partially induced in *Sprn*-knockdown, *Prnp*^*0/0*^ embryos, resulting in a high rate of embryonic lethality [2].

## 3.5 Conclusion

Overall, our results suggest a genetic adaptation of *Prnp*^*0/0*^, *Sprn*^*0/0*^ mouse embryos, both to sustain placental physiology that is affected in the absence of PrP [29] or Shadoo [7] and to refrain the upregulation of induced interferon responses following environmental stresses. This genetic adaptation might involve the downregulation of *Ada* and its-related pathways, this protein being involved in immunomodulation and ectoplacental development. Although this hypothesis remains to be further supported by direct experiments, it offers an explanation for the discrepancy observed between knockdown and knockout in previously reported data [2,3] and adds to the list of knockout genotypes that have acquired genetic adaptation.

## Funding

This work was supported by an INRAE Animal Genetics department project grant (grant AAP.GA.2015, COSI-Net).

## Notes

https://data.inrae.fr/privateurl.xhtml?token=e7b8885d-9cef-43c9-ae98-06259ee84d44

### Competing Interest Statement

The authors have declared no competing interest.

